# Illuminating Uveitis: Metagenomic Deep Sequencing Identifies Common and Rare Pathogens

**DOI:** 10.1101/054148

**Authors:** Thuy Doan, Michael R. Wilson, Emily D. Crawford, Eric D. Chow, Lillian M. Khan, Kristeene A Knopp, Dongxiang Xia, Jill K. Hacker, Jay M. Stewart, John A. Gonzales, Nisha R. Acharya, Joseph L. DeRisi

## Abstract

**Background:** Ocular infections remain a major cause of blindness and morbidity worldwide. While prognosis is dependent on the timing and accuracy of diagnosis, the etiology remains elusive in ~ 50% of presumed infectious uveitis cases.^1,2^ We aimed to determine if unbiased metagenomic deep sequencing (MDS) can accurately detect pathogens in intraocular fluid samples of patients with uveitis.

**Methods:** This is a proof-of-concept study, in which intraocular fluid samples were obtained from 5 subjects with known diagnoses, and one subject with bilateral chronic uveitis without a known etiology. Samples were subjected to MDS, and results were compared with conventional diagnostic tests. Pathogens were identified using a rapid computational pipeline to analyze the non-host sequences obtained from MDS.

**Findings:** Unbiased MDS of intraocular fluid produced results concordant with known diagnoses in subjects with (n=4) and without (n=1) uveitis. Rubella virus (RV) was identified in one case of chronic bilateral idiopathic uveitis. The subject’s strain was most closely related to a German RV strain isolated in 1992, one year before he developed a fever and rash while living in Germany.

**Interpretation:** MDS can identify fungi, parasites, and DNA and RNA viruses in minute volumes of intraocular fluid samples. The identification of chronic intraocular RV infection highlights the eye’s role as a long-term pathogen reservoir, which has implications for virus eradication and emerging global epidemics.

## INTRODUCTION

Ocular infection is an important cause of ocular morbidity and blindness worldwide. However, diagnosis is challenging due to the multitude of possible pathogens. Sensitivity of culture-based assays ranges from 40-70%, and available molecular diagnostics target only a fraction of pathogens known to cause ocular disease.^1–3^ These limitations are exacerbated by: (1) the inability to collect large intraocular fluid volumes given the eye’s small and delicate anatomy, and (2) the difficulty distinguishing clinically between infectious and non-infectious causes of ocular inflammation.

The urgency to develop better diagnostics for uveitis has been compounded by the recent cases of persistent infection with Ebola virus^4^, and possibly Zika virus.^5^ These cases highlight the eye’s role as a potential reservoir for infectious agents with important public health consequences. It is essential that more sensitive, unbiased, and comprehensive approaches are developed to efficiently diagnose ocular infections.

Rapid advances in sequencing technology and bioinformatics have made metagenomics a fertile area for developing clinical diagnostics.^6–8^ We evaluated the utility of an hypothesis-free approach to identify ocular infections by performing unbiased metagenomic deep sequencing (MDS) on total RNA extracted from the intraocular fluid of subjects with inflammatory and non-inflammatory eye diseases.

## METHODS

### Study Design

Six subjects were recruited for a research study using unbiased MDS to identify potential pathogens in intraocular fluid (aqueous or vitreous) (Table 1). This study was approved by the Institutional Review Board of the University of California, San Francisco (UCSF). Five of the six subjects served as controls to benchmark the ability of MDS to identify a variety of pathogens; Subjects 1-3 had ocular infections with herpes simplex virus 1 (HSV-1), *Cryptococcus neoformans*, and *Toxoplasma gondii*, respectively. Subject 4 had non-infectious uveitis, and subject 5 had no ocular inflammation but had intraocular fluid obtained at the time of a retinal membrane peel. MDS was also used to investigate subject 6 who had bilateral uveitis that had defied a 16-year diagnostic work-up at multiple academic centers across two continents (Table 1 and Figure 1A).

**Table 1:**
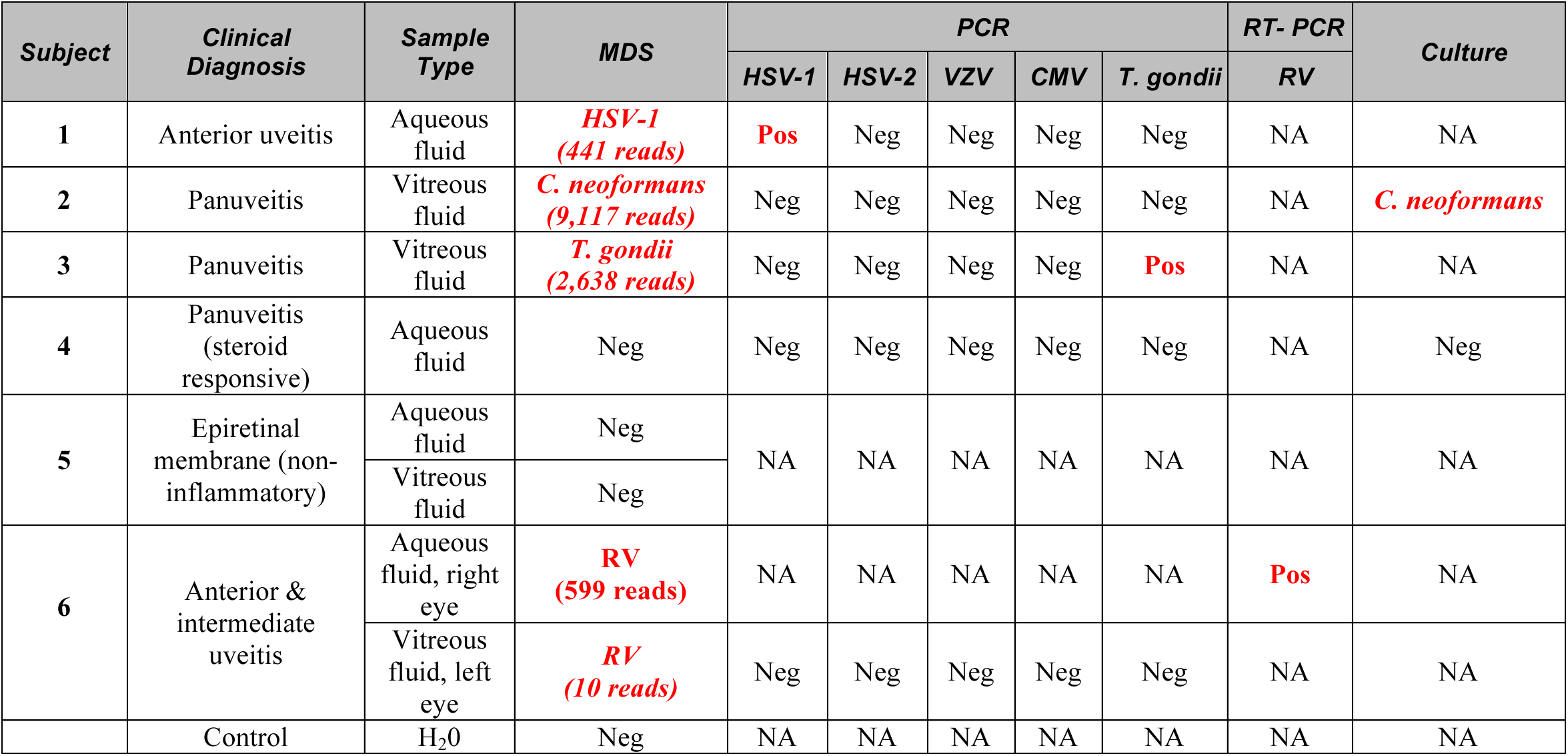
Results of Unbiased Metagenomic Deep Sequencing (MDS) and Conventional Diagnostic Tests on Intraocular Fluid Samples. MDS correctly identifies known infections in subjects 1-3. Subjects 4 and 5 had non-infectious ocular disease and had negative MDS testing for pathogens, defined as the presence of no microbial sequences other than known laboratory and environmental contaminants. Rubella virus was identified via MDS in subject 6 and confirmed by the California Department of Public Heath’s RT-PCR assay. Abbreviations: Pos, positive; Neg, negative; NA, not applicable; RT-PCR, reverse transcription polymerase chain reaction; HSV-1, herpes simplex virus-1; HSV-2, herpes simplex virus-2; VZV, varicella zoster virus; CMV, cytomegalovirus; *T. gondii, Toxoplasma gondii*; RV, rubella virus; *C. neoformans, Cryptococcus neoformans*; RE, right eye; LE, left eye.

**Figure 1:**
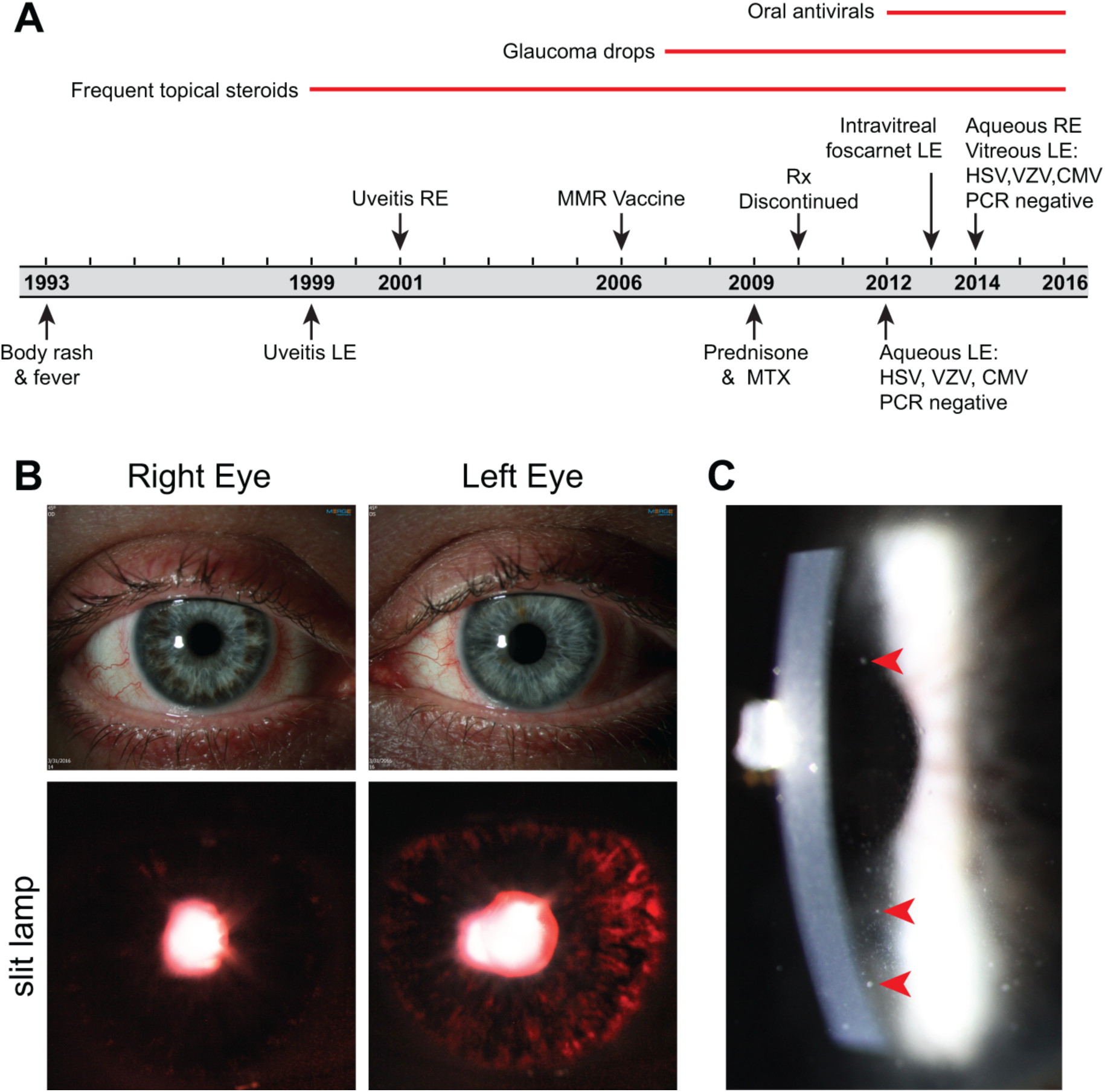
Clinical Course and Ocular Findings of a 40 Year-Old Man With Bilateral, Idiopathic Chronic Anterior and Intermediate Uveitis. Panel A shows Subject 6’s clinical course spanning 22 years. Panel B shows different colored irises (heterochromia) between the right and left eyes (top panels) and transillumination defects that are prominent in the left eye because of iris atrophy (lower panels). Panel C shows diffused aggregates of inflammatory cells (keratic precipitates; red arrows) on the endothelium of the cornea. Abbreviations: HSV, herpes simplex virus; VZV varicella zoster virus; CMV, cytomegalovirus; PCR, polymerase chain reaction; RE, right eye; LE, left eye; MMR, measles/mumps/rubella vaccine; MTX, methotrexate.

### Sequencing Library Preparation

Samples were prepared for MDS as previously described.^6^ Briefly, RNA was extracted from 20-50 μL of intraocular fluid and randomly amplified to double-stranded complementary DNA (cDNA) using the NuGEN Ovation v.2 kit (NuGEN, CA). cDNA was tagmented with Nextera (Illumina, CA). <u>D</u>epletion of <u>A</u>bundant <u>S</u>equences by <u>H</u>ybridization (DASH), a novel molecular technique using the CRISPR (Clustered Regularly Interspaced Short Palindromic Repeats)-associated nuclease Cas9 *in vitro*, selectively depleted human mitochondrial cDNAs from the tagmented library, thus, enriching the MDS library for non-human (i.e., microbial) sequences.^9^ One sample was prepared with New England Biolabs’ (NEB) Next modules to generate cDNA and the NEB Next Ultra II DNA kit to convert the cDNA into sequencing libraries (NEB, MA). Library size and concentration were determined using the Blue Pippin (Sage Science, MA) and Kapa Universal quantitative PCR kit (Kapa Biosystems, Woburn, MA), respectively. Samples were sequenced on an Illumina HiSeq 2500 instrument using 135/135 base pair (bp) paired-end sequencing.^6,7^

### Bioinformatics

Sequencing data were analyzed using a rapid computational pipeline developed by the DeRisi Laboratory to classify MDS reads and identify potential pathogens by comparison to the entire National Center for Biotechnology Information (NCBI) nucleotide (nt) reference database.^6^ The full dataset for each subject was analyzed in less than five minutes. Briefly, paired-end reads were quality filtered using PriceSeqFilter.^10^ Human sequence was removed by alignment to the human reference genome (hg38) using STAR.^11^ Unaligned reads that were at least 95% identical were compressed by cd-hit-dup (v4.6.1). These reads were then used as queries to search the NCBI nt database (July 2015) using gsnapl (v2015-12-31).^12^

## RESULTS

### MDS to Detect Pathogens in Uveitis

MDS accurately detected viral (HSV-1), protozoan (*T. gondii*), and fungal (*C. neoformans*) infections in subjects 1-3 and did not detect microbes other than known laboratory and environmental contaminants in subjects 4 and 5 (Table 1).

In subject 6, MDS detected a single candidate pathogen: rubella virus (RV) in an aqueous fluid specimen collected in 2014. 599 non-redundant sequence pairs mapped to both the non-structural and structural open reading frames (ORFs) of the RV genome. No sequences aligning to RV were present in the water control or the 18 other cerebrospinal or intraocular fluid samples sequenced on the same run. No RV reads have ever been detected previously in this laboratory.

Subject 6 was a 40 year-old man with a 16-year history of inflammation in both eyes whose extensive diagnostic work-up in Germany and the U.S. had been unrevealing (Table 1 and Figure 1A). In 1993 he had a three-day febrile illness accompanied by a rash that spread from his back to his extremities. He was diagnosed with anterior uveitis of the left eye in 1999 and in 2001, developed anterior uveitis of the contralateral eye. Topical steroid and non-steroidal anti-inflammatory drops were ineffective. Oral steroids were added in 2009 followed by methotrexate. His inflammation did not improve after one year of combined immunotherapy, and his medications were discontinued.

He presented to the Francis I. Proctor Foundation and UCSF in 2012 with moderate anterior and intermediate uveitis associated with ocular hypertension and diffuse stellate keratic precipitates in both eyes (Figure 1C) and asymmetrical iris atrophy leading to heterochromia (Figure 1B). These findings were suggestive of viral-related uveitis, and the subject underwent an anterior chamber paracentesis of the left eye. 100 μL of aqueous fluid was sent for polymerase chain reaction (PCR) testing for cytomegalovirus (CMV), varicella-zoster virus (VZV), and HSV-1/2. Despite negative results, suspicion for viral infection remained high. Antiviral therapy was initiated and continued for three years (Figure 1A), but failed to curb the inflammation. In 2014 he had a paracentesis of the right eye and a therapeutic vitrectomy of the left eye. Repeat infectious disease diagnostics were unrevealing (Figure 1A).

### Confirmatory testing for RV infection

A 183 nt RNA fragment was reverse transcribed and amplified from the subject’s aqueous fluid collected from the right eye in 2014, using a published reverse transcription PCR (RT-PCR) assay for detecting the RV E1 gene.^13^ Sanger sequencing confirmed that the amplicon was the RV E1 gene (Elim Bio, CA). This result was corroborated by the California Department of Public Health’s (CDPH) Viral and Rickettsial Disease Laboratory who performed RT-PCR and Sanger sequenced the 739 nt RV sequence required for genotype assignment (Sequetech Corp, CA).^14,15^ RV was not detected via RT-PCR in nasopharyngeal swab, urine, or tear samples collected in February 2016, indicating that the subject was not actively shedding virus. Serologic testing for RV IgG was positive.

An archived sample from the subject’s 2014 left eye vitrectomy subsequently underwent MDS using the same protocol. While the sample was not flash-frozen and was not stored to optimally preserve RNA integrity, ten unique sequence pairs aligned to the RV non-structural ORF. The detection of RV in both eyes corroborated the clinical suspicion of bilateral viral infection and demonstrated the robustness of MDS to detect pathogens despite suboptimal sample handling.

### Characterization of RV Sequences

The subject’s original MDS data were combined with sequencing data obtained from four replicate sequencing runs. These reads were aligned using bowtie2 v2.2.8 to the complete RV genome (GenBank DQ388280.1).^16^ 9,188 bp mapped, covering 95.1% of the reference genome (Figure 2A). This represents the most extensive coverage of an RV genome detected from any intraocular sample and suggests that the RV genomes are full length.^17^

**Figure 2:**
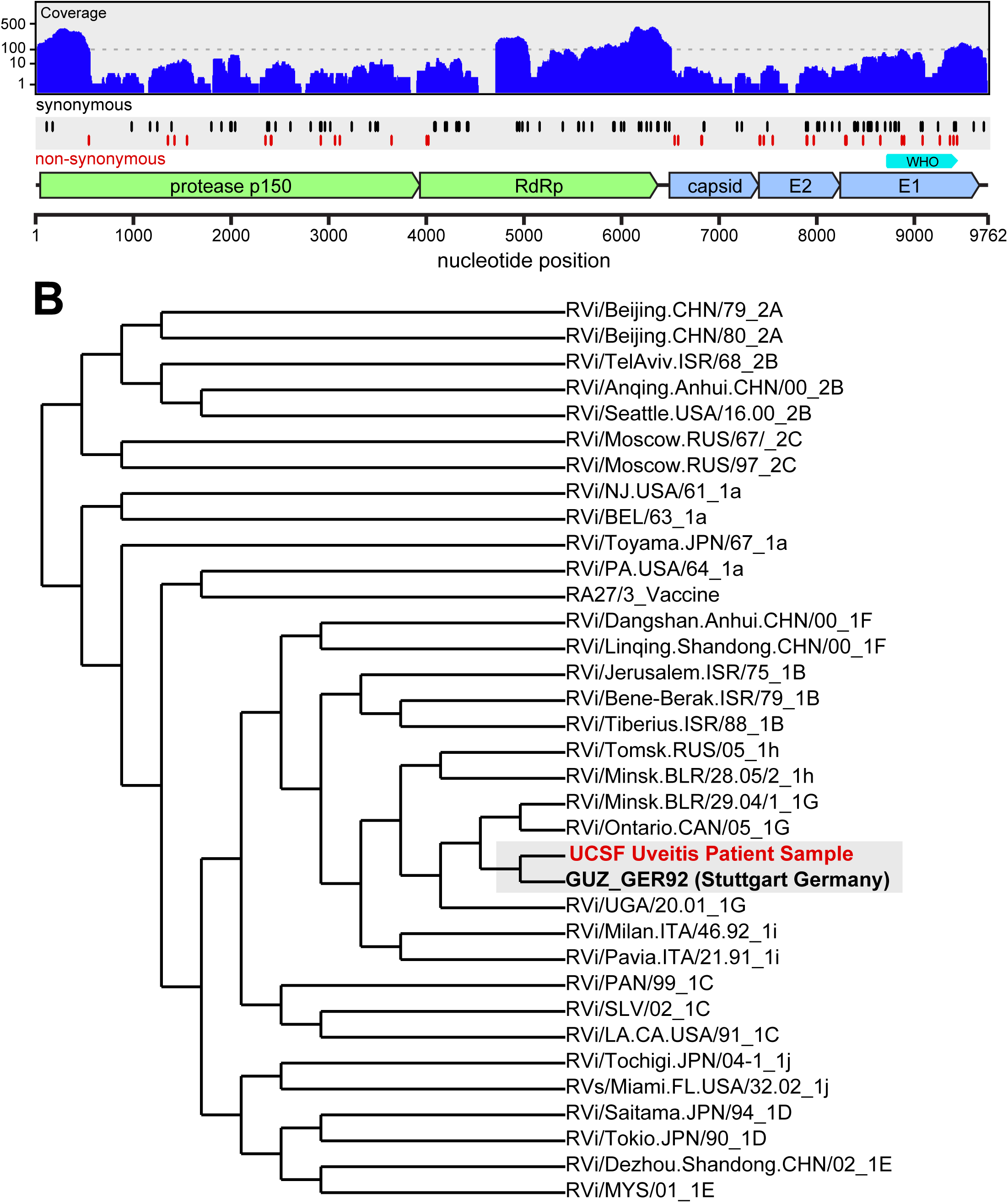
Identification of Rubella Virus (RV) by Metagenomic Deep Sequencing. Panel A illustrates how the 9,188 nucleotide (nt) paired-end sequence reads obtained from sequencing the RNA extracted from the subject’s aqueous fluid aligned to the most closely matched RV genome (DQ388280.1). 95.1% of the total RV genome is represented. Positions of synonymous (black vertical lines) and non-synonymous (red vertical lines) variants are shown. Of the 128 substitutions, 92 were synonymous, and 36 were non-synonymous. Of the 36 non-synonymous mutations, 19 occurred within the coding region for the E1 and E2 glycoproteins. Per unit length, the number of non-synonymous mutations in the E1 and E2 proteins was 4.1-fold higher than the non-structural proteins. The cyan marker above the E1 gene represents the 739 nt sequence window recommended by the World Health Organization (WHO) for RV genotyping. Panel B is a phylogenetic analysis of the subject’s RV strain obtained from MDS with 32 WHO reference strains, GUZ_GER92 (Stuttgart strain), and the RV27/3 vaccine strain, demonstrating that the subject’s RV sequence was most closely related to the genotype 1G viruses and not the vaccine strain.

### Phylogenetic analysis of the subject’s RV genome

The 739 nt segment of the RV E1 gene isolated from subject 6 with MDS was compared against the 32 World Health Organization (WHO) RV reference strains using <u>MU</u>ltiple <u>S</u>equence <u>C</u>omparison by <u>L</u>og-<u>E</u>xpectation (MUSCLE).^18–20^ His strain most closely aligned to the 1G genotype (Figure 2B). Of the three lineages of the 1G genotype, the lineage containing the Stuttgart strain circulated in Germany, Italy, and the United Kingdom in the early 1990s. Thus, this subject’s RV strain is temporally and geographically most proximate to the RV strain that was known to be circulating when he developed a rash and fever in 1993 in Germany.

The RV sequence (9,188 nt) obtained from our subject includes 128 nt substitutions relative to the 1992 Stuttgart strain (GenBank DQ388280.1). This substitution rate of 6.97 × 10^−4^ substitutions/site/year over the 20-year period is within two-fold of the RV evolutionary rate calculated as part of epidemiologic studies evaluating person-to-person transmission (1.19 × 10^−3^ to 1.94 × 10^−3^ substitutions/site/year).^21^ Of the 128 substitutions, 92 were synonymous (Figure 2A). Of the 36 non-synonymous mutations, 19 occurred within the coding region for the E1 and E2 glycoproteins. Per unit length, the number of non-synonymous mutations in the E1 and E2 structural proteins was 4.1-fold higher than the non-structural proteins. Considering all mutations in this region, the substitution rate in E1 and E2 was 1.05 x 10^−3^ substitutions/site/year. We note that this mutational imbalance associated with E1 and E2 compared to the non-structural proteins is consistent with persistent viral replication under immunological pressure.^22^

## DISCUSSION

Our results demonstrate that unbiased MDS can detect fungi, parasites, DNA viruses and RNA viruses in minute volumes of intraocular fluid from patients with uveitis. In addition to correctly identifying the causative agent in three infected positive control subjects (1-3) and detecting only background microbial contamination in two uninfected subjects (4 and 5), MDS revealed RV in a subject (6) who had a 16-year history of bilateral uveitis.

RV is a positive sense single-stranded RNA virus in the genus *Rubivirus* of the *Togaviridae* family that causes transient body rash and fever in healthy adults but can also cause devastating birth defects.^23^ RV has also been associated with Fuchs uveitis syndrome (FUS), a rare form of chronic intraocular inflammation most often characterized by mild anterior chamber reaction, iris atrophy with or without heterochromia, late onset ocular hypertension, and minimal associated visual complaints.^17,24,25^ In a subset of FUS patients, either RV IgG or RV RNA has been detected in ocular fluid by Goldmann-Witmer coefficient analysis or RT-PCR, respectively.^17,24,26^ These tests are only validated for ocular fluid at a few centers in Europe and are not diagnostically available in the U.S.

The protracted diagnostic challenge in our subject is three-fold: (1) diagnostic tests are lacking for ocular inflammation, (2) the subject’s clinical findings were not consistent with FUS until many years after disease onset, and (3) the subject’s relevant infectious exposure occurred six years prior to the onset of his ocular symptoms. This case highlights the advantage of an hypothesis-free approach in which a single MDS assay can detect a multitude of pathogens that may or may not have been previously associated with a particular clinical syndrome.

The identification of RV RNA in our subject’s eyes underscores current challenges in infectious disease surveillance. The WHO declared RV eliminated in the U.S. in 2005 as a result of effective and long-standing vaccination policies, but RV remains a threat throughout much of the world.^27,28^ Our subject’s ocular inflammation predated his measles, mumps and rubella (MMR) vaccination by seven years, and his RV strain most closely matched the strain circulating in his home country of Germany at the time of his rash and fever in 1993, and not the vaccine strain (Figure 2B). This is consistent with the notion that RV likely seeded his eyes during this primary infection. Although his immune system cleared the infection peripherally, RV sequestered in the ocular compartment and persisted presumably due to relative immune privilege. Indeed, our analysis of the RV genome provides the first molecular evidence for active RV replication in FUS. Ocular RNA virus sequestration is not a phenomenon relating solely to RV, as Ebola virus was recently detected in the ocular fluid of a patient nine weeks after resolution of his viremia.^4^ Using RT-PCR for RV on our subject’s tears, we were not able to detect shedding of RV, although longitudinal studies are required to determine whether intermittent shedding through tears can occur. As we devise strategies to rapidly identify and control emerging and re-emerging infectious diseases, expanding the scope of pathogen detection to the eyes and other immune privileged sites may be of critical importance.

Diagnostic tests for intraocular infection fundamentally differ from those for systemic infections because of the small sample volume that can be safely obtained from the eye. Unbiased MDS may circumvent this limitation, as it detects many infectious organisms with a single assay requiring as little as 20 μL of intraocular fluid. Not only does MDS have the potential to alter the paradigm for infectious disease diagnostics in ophthalmology, but it may also provide another valuable public health tool to surveil for re-emerging and emerging infectious diseases in immune privileged body sites.

## CONTRIBUTORS

TD and MRW contributed equally and therefore are co-first authors. JLD and NRA conceived the study. JLD, MRW, and TD developed study protocol and design, and were responsible for the study implementation and project management. TD, MRW, LMK, Emily D. Crawford, and Eric D. Chow performed library preparation and sequencing. JLD, MRW, and TD performed statistical analysis. TD, MRW, and KAK performed rubella RT-PCR. DX and JKH supervised the confirmatory rubella RT-PCR at the CDPH. TD, NRA, JG, and JMS obtained clinical samples and participated in patient care. TD, MRW, and JLD wrote the first draft of the article. All authors contributed to the interpretation of the data and the writing and editing of the article.

## DECLARATION OF INTEREST

We declare no competing interest.

## ACKNOWLEDGMENTS

Research reported in this publication was supported by a grant from the UCSF Resource Allocation Program for Junior Investigators in Basic and Clinical/Translation Science (T.D.); UCSF Center for Next-Gen Precision Medicine supported by the Sandler and William K. Bowes, Jr. Foundations (J.L.D. and M.R.W.); Howard Hughes Medical Institute (J.L.D.); the National Center for Advancing Translational Sciences of the NIH under Award Number KL2TR000143 (M.R.W.); the Cooperative Agreement Number U60OE000103, funded by the Centers for Disease Control (CDC) and Prevention through the Association of Public Health Laboratories (D.X. and J.K.H). Its contents are solely the responsibility of the authors and do not necessarily represent the official views of the CDC, the Department of Health and Human Services, the Association of Public Health Laboratories, or the NIH.

We thank Derek Bogdanoff in the UCSF Center for Advanced Technology for his expertise and assistance operating the Illumina sequencer and Dr. Steven Miller, Director of the UCSF Microbiology Laboratory, for his assistance coordinating confirmatory laboratory studies. We thank Daniela Munafo and Erbay Yigit from New England Biolabs with assistance with the sequencing library preparation. We thank the Measles, Mumps, Rubella & Herpesviruses Laboratory Branch at the CDC, particularly Emily Abernathy and Dr. Joseph Icenogle, for helpful discussions regarding the possible public health implications of the RV case. We thank the Sandler and William K. Bowes, Jr. Foundations for their generous philanthropic support. Lastly, we thank our patients for their participation in this research program.

